# Enrichment and delivery of target proteins into the cell cytosol *via* Outer Membrane Vesicles

**DOI:** 10.1101/2023.03.02.530906

**Authors:** Huan Wan, Zhiqing Tao, XiaoLing Zhao, Guan Wang, Yihao Chen, Juan Zhang, Xu Zhang, Maili Liu, Guosheng Jiang, Lichun He

## Abstract

Advanced intracellular delivery of proteins has profound applications in both scientific investigations and therapies. However, existing strategies relying on various chemical and physical methods, have drawbacks such as the requirement of high concentration *in vitro* prepared target proteins and difficulty in labeling target proteins. Developing new delivery systems integrating the enveloping and labeling of target proteins would bring great advantages for efficient protein transfections. Here, we enriched a high concentration (62 mg/ml) of several target proteins into outer membrane vesicles (OMVs) of *E. coli* to employ the native property of OMVs to deliver proteins into the cytosol of eukaryotic cells. The results revealed a high protein transfection efficiency arranging from 90-97% for different cell lines. Moreover, the free penetration of molecules less than 600 Dalton across the membrane of OMVs allows direct labeling of target proteins within OMVs, facilitating the visualization of target proteins. Importantly, the nanobody delivered intracellularly by OMVs retains the biological activity of binding with its target, highlighting the advantages of OMVs as an emerging tool for efficient intracellular delivery of proteins.

## Introduction

Intracellular delivery of proteins has broad applications in both scientific research and medical therapies.^1^ Important applications include the delivery of labeled cargo proteins for cell imaging, diagnosis, and signaling pathway research. For example, direct tagging of the interested cargo protein with a fluorescent molecule allows tracking the transportation and localization of proteins with a high spatial resolution in cells *via* fluorescence microscopy.^2-5^ Likewise, new medicines based on proteins are becoming more and more appealing for drug development. However, most of the targets of the protein drugs are restricted to the extracellular receptors due to the limitation of available proper cytosolic delivery approaches. The efficient protection and transportation of the proteins into the cell could largely expand targets of human diseases, providing more choices to use antibodies for the medical treatment of numerous diseases.^6^

As the significance of intracellular protein delivery has been well recognized, multiple methods were developed for cellular protein delivery including physical methods, cell-penetrating peptides, polyethyleneimine, supercharged molecules, and nanocarriers in the past decades.^7-11^ Most of these transportation systems have remained limitations in different aspects. Physical methods such as electroporation, microinjection, or microinjection, mainly use physical forces to temporally generate pores on the cell membrane to allow proteins to transport into cells, but they often induce damage to cell membranes and have limited throughput.^12^ Approaches of using cell-penetrating peptides, polyethyleneimine, and supercharged molecules usually face problems of poor delivery efficiency due to cargo proteins weakly interacting with these transportation systems. Similarly, nanoparticles including lipid-based nanoparticles, polymer micelle nanoparticles, and inorganic nanomaterials, mainly bind with the cargo proteins noncovalently.^13, 14^ Therefore, they need to be customized according to different surface characters of cargo proteins for high transportation efficiency. Toward overcoming deficiencies of aforementioned delivery methods, we aim to develop a new system, which could envelop and enrich various cargo proteins regardless of the binding efficiency of cargo proteins with the delivery system. Outer membrane vesicles (OMVs) are extracellular vesicles released by bacteria into the extracellular milieu to interact with host cells to establish bacteria-host mutualism.^15^ The diameter of OMVs ranges from 50-250 nm, packing various biomolecules in their lumen spaces. Previous reports revealed OMVs exhibited extremely high biocompatibility with host cells as their composition was similar to cell membranes.^16, 17^ Hence it is promising to modify this OMV-mediated biomolecules transport pathway for delivering biologically active proteins into the target cells.^18^

In this work, we demonstrated the feasibility of directly enriching and labeling target proteins in the lumen of OMVs. Moreover, OMVs enabled the enveloped proteins including the nanobody to directly penetrate across the cell membrane and remain in the active state, strongly suggesting that intracellular protein delivery *via* OMVs will have genuine utility in fields of biology research and therapeutics.

## Results and Discussion

### Characterization of OMVs

OMVs were produced and purified from the cell culture of *E. coli* according to a published protocol.^19^ A final filtration of OMVs through a 0.22 μm cellulose filter membrane was applied to polish and sterilize the OMVs. With the OMVs obtained, transmission electron microscopy (TEM) and nanoparticle tracking analysis (NTA) were performed to characterize the surface properties of OMVs. TEM images revealed the produced OMVs had round morphologies with clear edges, suggesting complete and intact membranes were present (Figure 1a). NTA results of OMVs showed a narrow range of size distribution from 50 to 200 nm, with an average size of 110 nm (Figure 1b) and a negative zeta potential with the value of -40.49 ± 0.76 mV, indicating good stability of the OMVs suspensions.

**Figure 1.**
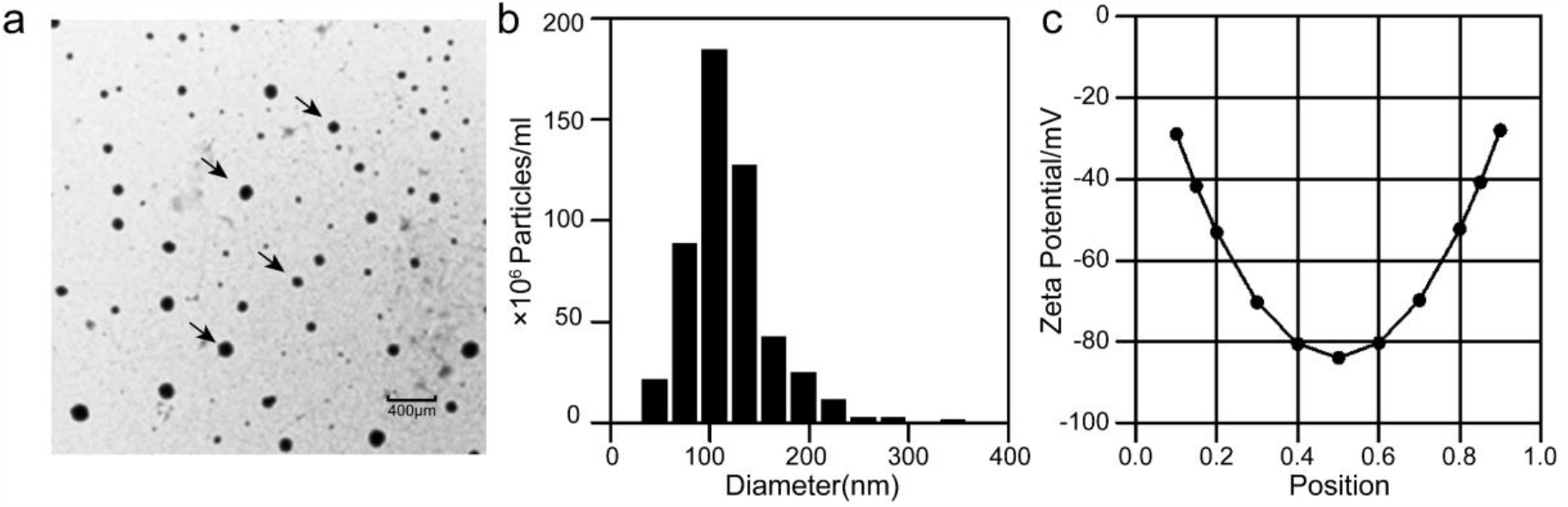
Properties of purified OMVs. (a) TEM images of OMVs (scale bar, 200nm). (b-c). Particle size distribution (b) and Zeta Potential (c) of OMVs as determined by NTA

### Protein-delivery capacity and cytotoxicity of OMVs

As OMVs are naturally evolved vehicles for intraspecies and interspecies biomolecule communication, including nucleic acid and proteins. We hypothesized that the enrichment of target proteins in the lumen of OMVs would be an attractive way for intracellular protein delivery in at least two aspects: the low cost of producing the OMVs by culturing *E. coli* and directly labeling target protein in OMVs with various dyes.^20^ To test this, we first established a secretion system to translocate the overexpressed protein into the periplasm, facilitating its subsequent envelopment into the OMVs. A model protein Spy in our lab fusing with the pelB signal peptide at the N-terminus was cloned into the pET-21a vector (Figure 2a). The fused protein was then expressed as published before^21^ and analyzed by the sodium dodecyl sulfate-polyacrylamide gel electrophoresis (SDS-PAGE). Spy was successfully expressed and highly enriched into the OMVs as revealed in Figure 2c. To assess the concentration of Spy in OMVs, we first quantified intensities of SDS-PAGE gel bands of purified Spy samples with different concentrations to make a standard curve with the software ImageJ. Then the apparent concentration of Spy in the OMVs suspension was calculated based on this standard curve. The radius and numbers of OMVs from NTA measurements were used to calculate the total inner volume of OMVs. With the ratio of the volume of OMVs suspension and the total inner volume of OMVs, a final concentration of Spy in OMVs was estimated to be ∼62 mg/ml (Figure 2d), proving the high efficiency of loading target protein onto the delivery system. Moreover, the membrane of OMVs hosted many pores formed by outer membrane proteins. Some of the pores allowed molecules with molecular weights less than 600 Da to pass over, and show no particular substrate specificity.^22^ We then tested if we could label the enriched proteins with fluorescent dye directly. A cysteine mutant T72C of Spy was introduced by the site-directed mutagenesis and expressed in the same way in *E. coli*. The OMVs were collected and purified to incubate with Dylight 488 sulfhydryl-reactive dyes (Dylight 488) at 4 ℃ overnight. The free Dylight 488 was then removed with a 14 kDa cut-off dialyzes cassette (Figure 2b). Comparing with the Coomassie-stained SDS-PAGE gel of OMVs enriching Spy (Figure 2e left), the image of the same gel captured under an excitation light of 488 nm revealed a single clear band of labeled Spy (Figure 2e right), while bands of other proteins, mainly the membrane proteins on OMVs, were not visible, manifesting the successful labeling of the target Spy protein by Dylight 488 within OMVs.

**Figure 2.**
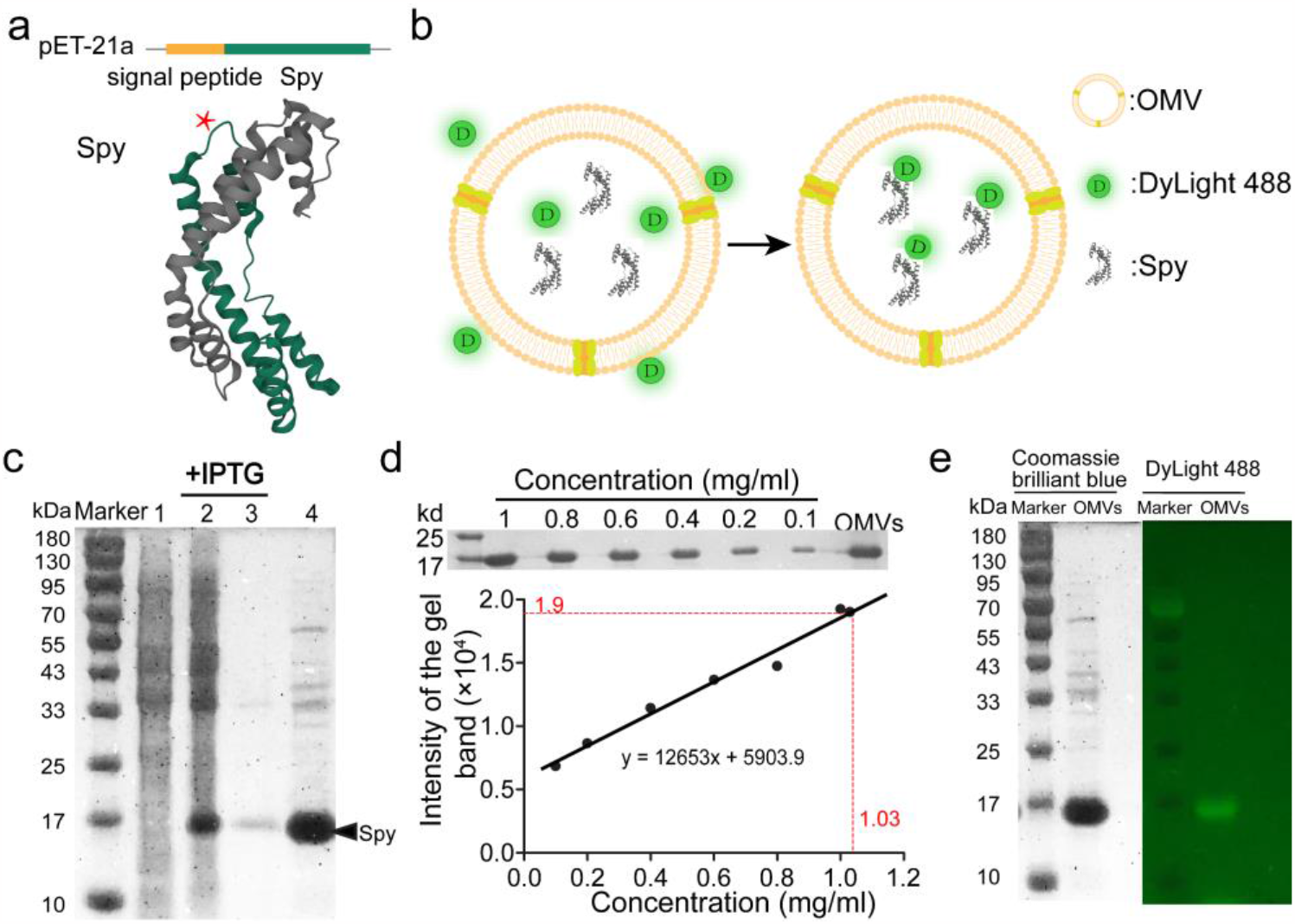
Enrichment and labeling of Spy in the lumen of OMVs. (a) Schematic diagram of the secreted expression plasmid. Target protein (Spy) fused with the pelB signal peptide at the N-terminus. The location of the introduced site mutagenesis T72C of Spy was indicated by * in the crystal structure of Spy (PDB code 3O39). (b) Schematic diagram of labeling Spy with DyLight 488™ Sulfhydryl-Reactive Dyes. (c) SDS-PAGE analysis of the enrichment of Spy in OMVs: lane 1, sample of the control cell pellet; lane 2, sample of the cell pellet; lane 3, sample of the culture medium; lane 4, sample of the harvest OMVs. The band of Spy was indicated with a black arrow. (d) SDS-PAGE gel band of OMVs enriched with an unknown amount of Spy and gel bands of the purified Spy with different concentrations (unit in mg/ml) (upper). The standard curve of the gel band intensity plotted against the concentration of Spy. The determined apparent concentration of Spy in the OMV sample was graphed in red (lower). (e) SDS-PAGE analysis of the directly labeled Spy sample by DyLight 488™ Sulfhydryl-Reactive Dyes within OMVs. Image of Coomassie brilliant blue stained SDS-PAGE gel of OMVs enriching the T72C mutant of Spy (left). Image of the SDS-PAGE gel of OMVs enriching the labeled T72C mutant Spy captured under a light with a wavelength of 488 nm (right).

### Protein-delivery capacity and cytotoxicity of OMVs

We next investigated whether OMVs could deliver their encapsulated proteins into eukaryotic cells. 75 µg sterile OMVs enriched with Dylight 488-labeled Spy were co-incubated with Hela, 293T, and CX-1 cells in 24 well plates for 12h respectively. The localization of labeled Spy protein in cells was tracked by a confocal Cell laser scanning microscopy (CLMS). Punctate green fluorescence was observed all over the cytoplasm of all three different cell lines (Figure 3), indicating the labeled Spy was successfully delivered into the cell. The transfection efficiencies of labeled Spy into Hela, 293T, and CX-1 cells were determined to be 92%, 90%, and 97% with the fluorescence microscopy respectively (Figure 4b). To access the cytotoxic effect of OMVs on target cells, we first compared the morphology of Hela, 293T, and CX-1 cells before and after the treatment with OMVs. No significant morphological changes were observed for all three cell lines after the incubation with OMVs for 16h (Figure 4a). Additionally, colorimetric assays with CCK-8 revealed both the proliferation and viability of the cells treated with OMVs increased similarly to the controlled OMV buffer treated group over 72 hours, confirming the negligible cytotoxicity of OMVs (Figure 4c-e).

**Figure 3.**
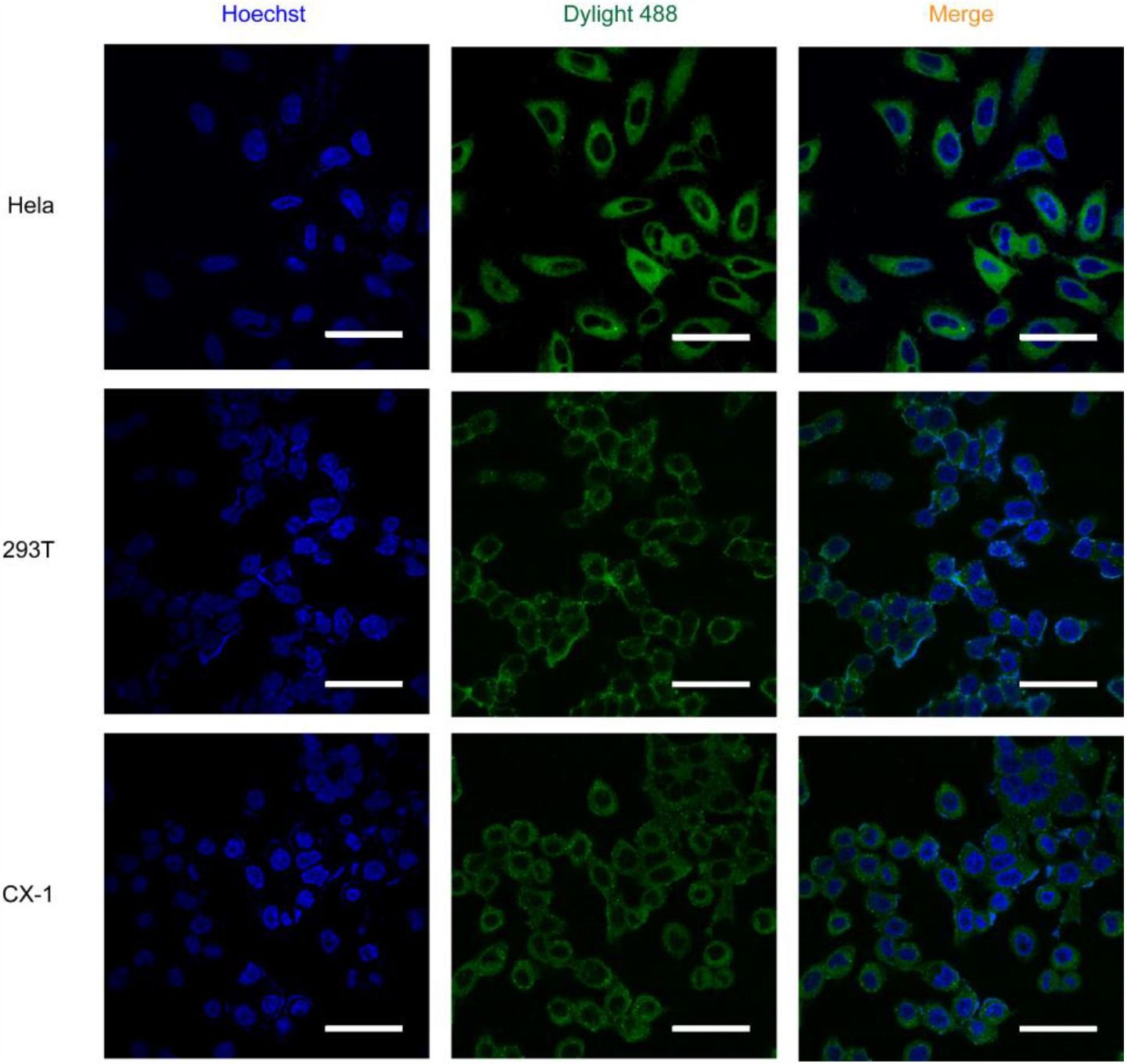
CLMS images of Hela, 293T, and CX-1 cells following 12h of incubation with OMVs enriching Dylight 488 labeled Spy (Scale bar, 50μm). Nuclei were stained with Hoechst dye (blue).

**Figure 4.**
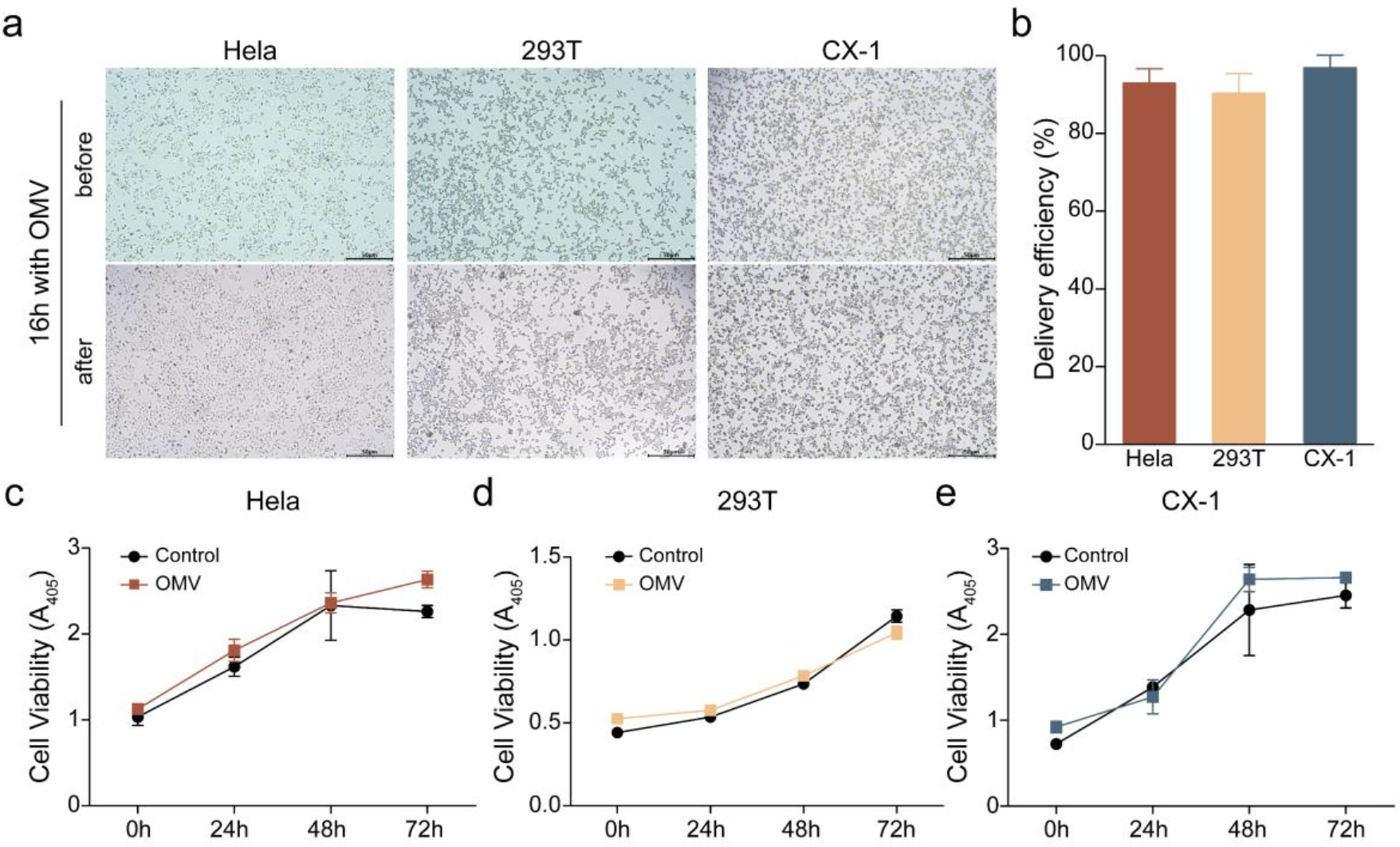
Verification of the OMVs delivery system on various cells. (a) Microscope images of Hela, 293T, CX-1 cells before and after incubation with OMVs for 16 h. (b) Quantification of the intracellular delivery efficiencies of labeled Spy protein into three different cell lines *via* OMVs over 24 h incubation (n=4). (c-d) viability of Hela (c), 293T (d), and CX-1 (e) cells. Cells treated with OMVs or the OMV buffer for different durations (0h, 24h, 48h, and 72h).

### Mechanism of the internalization of OMVs into cells

To further characterize the cell entryway of OMVs, we first labeled OMVs with a lipophilic membrane stain (1, 1’-dioctadecyl-3, 3, 3’, 3’-tetramethylindocarbocyanine perchlorate (Dil)). The labeled OMVs were then incubated with Hela, 293T, and CX-1 cell lines respectively. CLMS was applied to monitor the uptake of OMVs by different cell lines. Figure 5a-c revealed the internalization of OMVs by each cell line in a time-dependent way. The red fluorescence from Dil-labeled OMVs accumulated gradually in cytosol along with the incubation time. In addition, the amount of OMVs entering cells varied among different cell lines. The fluorescence intensity of Dil-labeled OMVs in CX-1 cells was weaker than the other two cell lines, indicating the number of OMVs internalized by each CX-1 cell was less (Figure 5a-c). It was worth noting that the high transfection efficiency of CX-1 determined in Figure 4b did not refer to the number of the OMVs or encapsulated proteins internalized, but the percentage of cells showing positive fluorescence.

**Figure 5.**
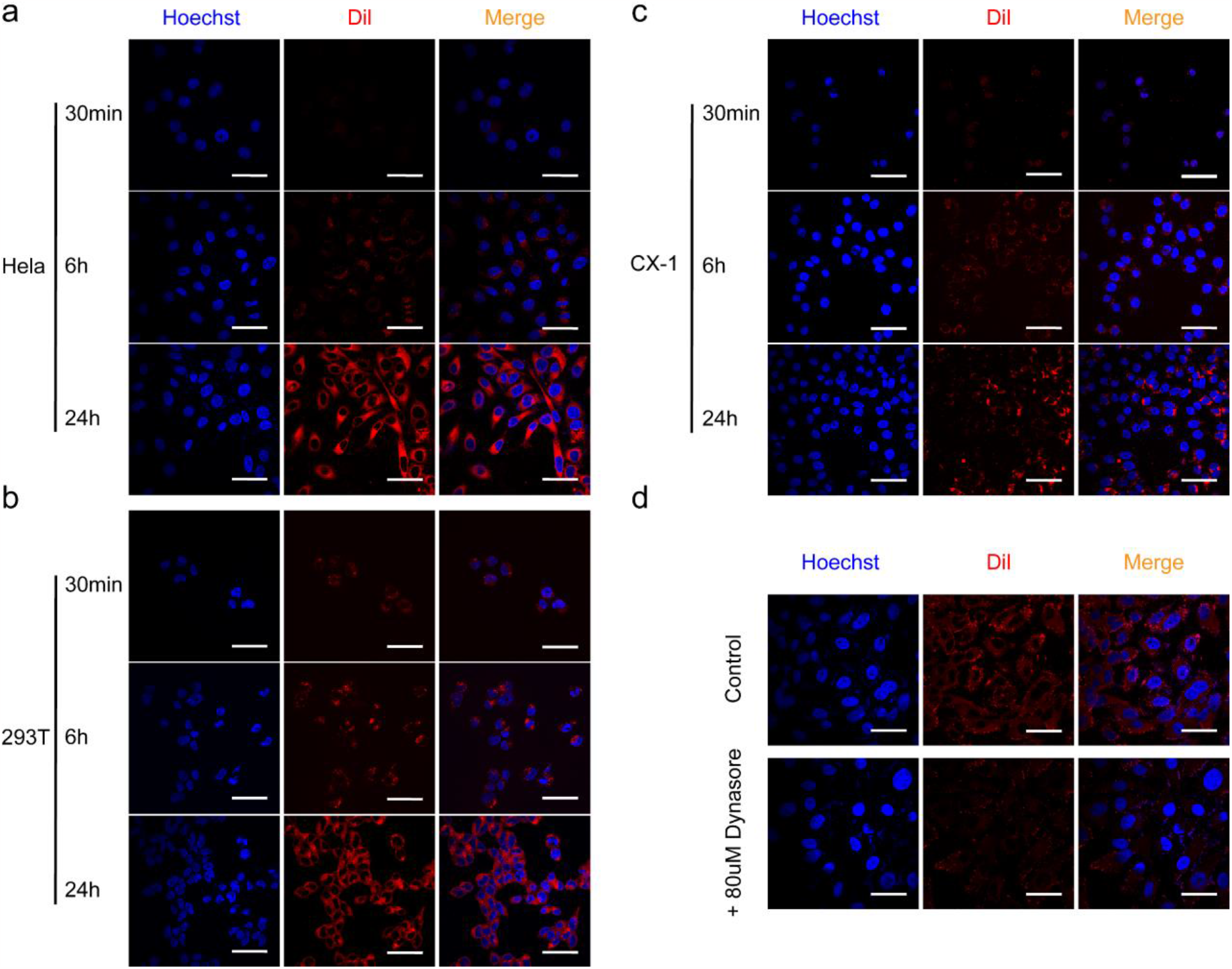
The internalization of OMVs by different cell lines monitored by CLMS. (a) CLMS images of Hela cells incubated with Dil-labeled OMV for 30min, 6h, and 24h (Scale bar, 50μm). (b) CLMS images of 293T cells incubated with Dil-labeled OMV for 30min, 6h, and 24h (Scale bar, 50μm). (c) CLMS images of CX-1 cells incubated with Dil-labeled OMV for 30min, 6h, and 24h (Scale bar, 50μm). (d) CLMS images of Hela cells incubated with Dil-labeled OMV for 6h with and without the pre-treatment of 80mM Dynasore (Scale bar, 50μm).

Notably, the punctate red fluorescence of Dil-labeled OMVs spread wildly in the cytoplasm, implying that OMVs may enter the cells by endocytic pathways. To confirm this, we analyzed the effects of an inhibitor of endocytosis, Dynasore, which was capable of interfering with the neuronal endocytic protein dynamin-1. Hela cells were pre-incubated with Dynasore in the cell culture medium at a concentration of 80 mM for 30 min, where the control group of Hela cells was pre-incubated with the same volume of OMV buffer. Figure 5d showed the fluorescence intensity of Dil-labeled OMVs in the cytosol of Hela cells treated with Dynasore was significantly weaker than the control group, indicating Dynasore was able to reduce the intracellular uptake of OMVs. This result suggested that dynamin-dependent endocytosis was essential for the internalization of OMVs.

### Intracellular delivery of nanobodies by OMVs

Last but not the least, we tested if the nanobody could be delivered intracellularly through the OMV system. An arbitrary nanobody cloned from the naïve library was constructed into the pelB secretion vector and overexpressed in *E. coli* as aforementioned. SDS-PAGE gel revealed the nanobody was also highly enriched in the OMVs. The labeling of nanobodies with Dylight 488 was carried out as before. The intercellular delivery of labeled nanobody into Hela, 293T, and CX-1 cells was confirmed by CLSM (Figure 6).

**Figure 6.**
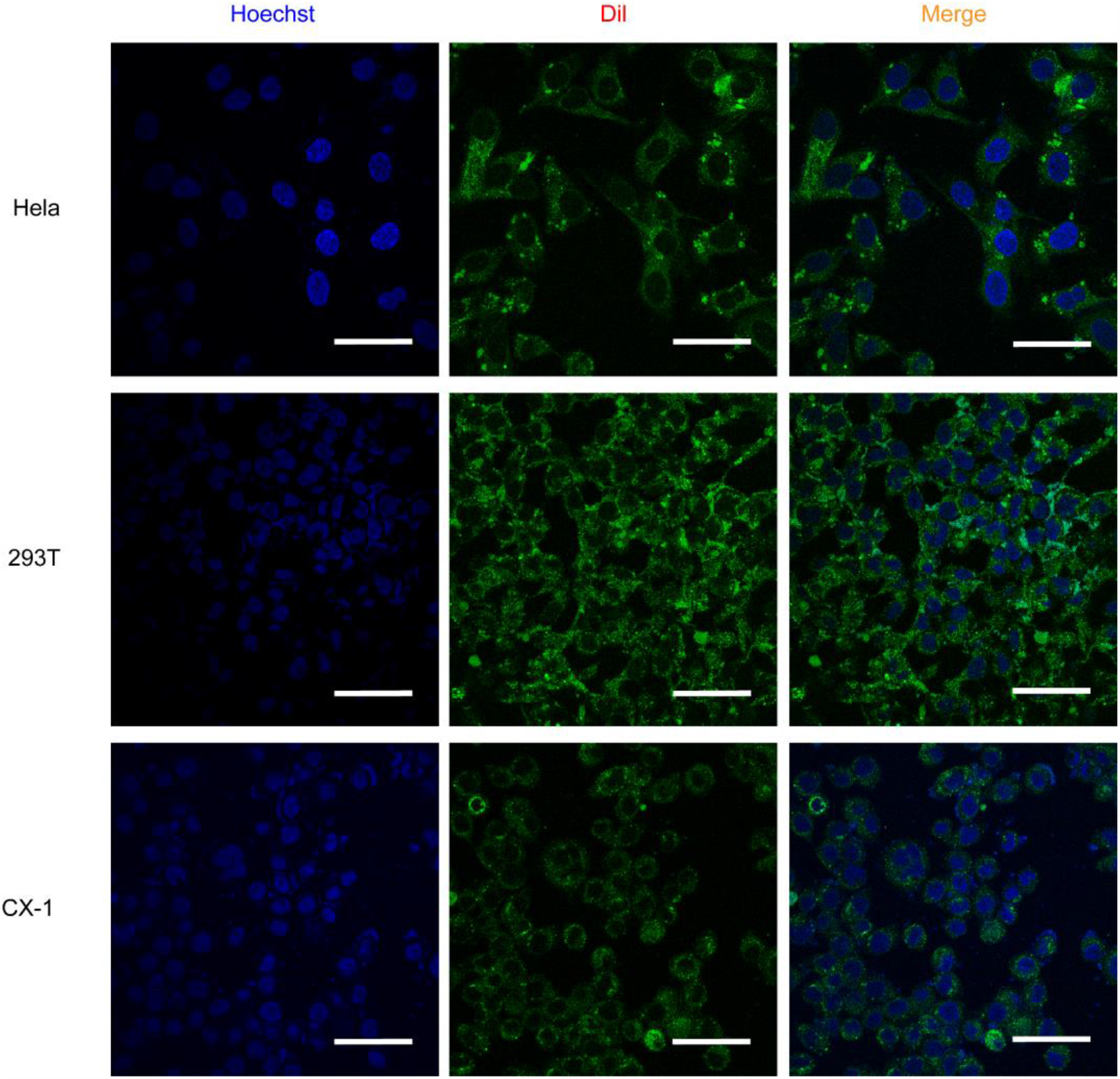
Intracellular delivery of the labeled nanobody into different cell lines by OMVs. CLSM images of Hela, 293T, and CX-1 cells following 12h of incubation with OMVs enriching DyLight 488 labeled nanobodies (Scale bar, 50μm).

To determine the biological activity of the internalized protein through OMVs, a Pfu DNA polymerase specific nanobody screened in our lab was expressed and enriched in OMVs. The Hela cells incubated with control OMVs and OMVs enriching the Pfu nanobody were collected and lysed respectively. The supernatants of cell lysates were applied for enzyme-linked immunosorbent assay (ELISA), where the plates were coated with 10 ng Pfu DNA polymerase per well. The OD_450_ value of the OMVs group enriching Pfu nanobody was similar to that of the *in vitro* purified Pfu nanobody group. This result demonstrated the internalized nanobody retained its activity of binding with the target protein, expanding the utility of the OMV system for direct labeling and transfections of nanobodies (Figure 7).

**Figure 7.**
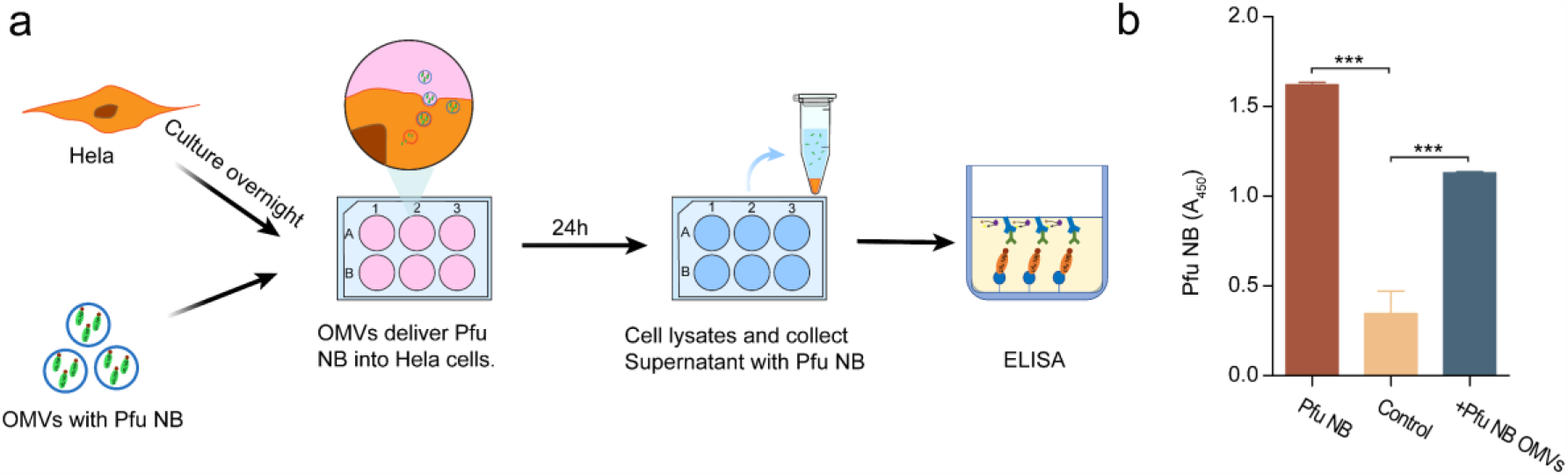
The bioactivity of the Pfu nanobody transfected into Hela cells *via* OMVs. (a) Scheme for validating the activity of the internalized Pfu nanobody (b) ELISA results of lysates of Hela cells incubated with a purified Pfu nanobody(10ng/ml), control OMVs and Pfu nanobody enriched OMVs. Results were indicated by OD_450_ using horseradish peroxidase (HRP) labeled secondary antibody as a tracer. Data were shown as means ± standard deviations from three independent experiments. *** p<0.001.

Cytosolic delivery of proteins is critical for direct cell manipulation either for protein therapeutics or for biological research.^23, 24^ Thus, developing an efficient intracellular protein delivery system would bring opportunities for new druggable intracellular targets and advanced investigations of cytosolic protein functions.^25^ The OMVs system described here is highly efficient and easily adaptable to assist the entering of intact biomolecules into the cytosol. By constructing into the pelB secretion expression vectors, various prokaryotic and eukaryotic target proteins are demonstrated to be enriched and enveloped into OMVs. The concentration of enriched cargo protein could reach ∼62 mg/ml (Figure 2c), avoiding lots of laborious work to get such a high concentration of purified cargo protein and to assemble it with the delivery system when compared to other protein delivery techniques. Furthermore, the cargo proteins in OMVs are still accessible for chemical modifications by organic dyes for real-time tracking and imaging of the cargo proteins in cells (Figure 2e). If desired, the modifications of cargo proteins with certain chemical groups such as positively charged groups could also assist the endosomal escape of the cargo proteins,^26^ allowing this OMV-based protein delivery system adjustable to endosomal stimuli. Thus, OMVs enriching cargo proteins could be envisioned for a broad scope of applications in basic biological research and translational medicines.^27^

With respect to biological studies, transfections of fluorescently labeled cargo proteins or nanobodies by OMVs will enable the direct visualization of locations and movements of target proteins. Instead of snapshots of continuous events of immobilized cells or tissues by conventional immunohistochemistry, the fully functional cycle of proteins is now trackable in real-time in living cells. Moreover, procedures for the treatment of samples are also much more simplified. With respect to translation medicines, protein drugs are still restricted due to their incapability to spontaneously penetrate cell membranes. Over 90% of the drugs on the market are small compounds as they could traverse the cell membrane to bind with intracellular targets.^28^ The OMVs-based cytosolic delivery approach developed in this work shows great potential to alter this situation. The capability of OMVs to deliver bioactive nanobodies into the cytosol offers possibilities to greatly expand antibody-based medical therapies to previously “undruggable” intracellular targets. Although the endotoxin from *E. coli* derived OMVs still hinders its direct application for *in vivo* treatment, developments and optimizations to produce endotoxin-free OMVs will not be far.^29-31^

## Conclusion

In summary, we extend the intraspecies and interspecies biomaterial transportation feasibility of OMVs for cytosolic delivery of cargo proteins. The developed OMVs system enables the enrichment of high concentrations (e.g., 62 mg/ml) of cargo proteins and allows direct labelling of cargo proteins within OMVs. This OMV-based approach brings new opportunities for fluorescent dye labeled proteins and therapeutic proteins to enter the cytoplasm, making it a promising method for simple and robust intracellular protein delivery.

## ASSOCIATED CONTENT

### Supporting Information

Experimental materials and methods (PDF)

### Author Contributions

Experiments were performed by Huan Wang, Zhiqing Tao, and Xiaoling Zhao; Data analysis and validation were done by Huan Wang, Zhiqing Tao, Xiaoling Zhao, Guan Wang, and Yihao Chen; The manuscript was written by Lichun He, Huan Wang, Zhiqing Tao, and Xiaoling Zhao; Experiments were designed by Lichun He, Guosheng Jiang, Xu Zhang, Juan Zhang, and Maili Liu. Funding was received by Lichun He and Maili Liu. All authors have approved the final version of the manuscript.

### Notes

The authors declare no conflict of interest in this work.

## ACKNOWLEDGMENT

This work was supported by the National Key R&D Program of China grants 2018YFE0202301 and 2018YFE0202300, National Natural Sciences Foundation of China grants 22174151, 21904138, and 21991080, Natural Science Foundation of Shandong grants ZR2022MH282, ZR2020QH160, ZR2021MH080.

