## Supporting Information for "Enrichment and delivery of target proteins into the cell cytosol *via* Outer Membrane Vesicles"

### Experimental

#### Isolations of Outer Membrane Vesicles (OMVs)

The sequence of Spy was obtained from the NCBI database. Gene of Spy was synthesized and cloned into pET-21a vector with N terminal fused with the pelB signal peptide by using FasHifi Super DNA polymerase (Swiss Affinibody LifeScience AG). Vectors of nanobodies were gifts from Qinhe Life Science Ltd, Wuhan, China. Plasmids were transformed into *E. coli* T7 competent cells, and then spread in ampicillin (Amp) resistant plates. A single clone was picked and incubated in 10ml LB-Amp medium for culturing overnight at 37°C as the preculture. 1L of M9 medium was then inoculated with the overnight preculture at 1:100 dilution, and induced with 0.5 mM Isopropyl-β-D-thiogalactopyranoside (IPTG, Aladdin) at 37°C for 15 h until the optical density at 600 nm ($OD_{600}$) reached 0.8. The culture medium was centrifuged twice at 5000 rpm at 4°C for 30 min. The supernatant was collected and filtered through 0.45 μm and 0.22 μm membrane filters successively for concentrating by Hydrosart® Microfilter (Sartorius). The concentrated supernatant was centrifuged at 20000 rpm at 4°C for 3 h to harvest the OMVs pellets, which were then carefully resuspended with phosphate-buffered saline (PBS).[1]

#### Characterization of OMVs

Transmission Electron Microscopy (TEM) was employed to measure the morphology of OMVs. The OMV preps were applied to a copper grid and blotted after soaking for 10 minutes and washed 3 times with dd$H_2O$. Samples were stained with 2% phosphotungstic acid for 3 minutes and washed 3 times by dd$H_2O$ and dried overnight. Photos of OMVs were taken on a TaLos L120C microscope operated at 120 kV. Hydrodynamics and Zeta potentials of isolated OMVs were assessed by Nanoparticles Tracking Analysis (NanoSight NS300).

#### Quantitative analysis of proteins in OMVs

Total protein concentration in OMVs was quantified with the bicinchoninic acid assay (BCA assay, Beyotime Biotecnology). To determine the concentration of the cargo protein in OMVs, gel bands of the cargo protein in OMVs were quantified with Image J and compared with a standard curve made with the purified cargo protein to determine its quantity. Vesicle radius and numbers of OMVs in a given $OD_{600}$ were measured with NTA to calculate the total inner volume of vesicles, which was then used to calculate the approximate concentration of the cargo protein in OMVs.

**Labeling of cargo proteins in OMVs**

OMVs enriching the T72C mutant of Spy were incubated with DyLight 488™ Sulfhydryl-Reactive Dye (Thermo Fisher) and Tris (2-carboxyethyl) phosphine (TCEP, Macklin) with a final concentration of 10 ug/ml and 10 mM at 4°C for 12 hours respectively. Excess dye reagent was removed by dialysis against PBS three times. The final sample of OMVs was filtered twice with a 0.22 μm filter for later usage. SDS-PAGE electrophoresis was used to visualize the labeled Spy within OMVs. In brief, OMVs were treated with 0.2% Triton X-100, followed by five freeze–thaw cycles to disrupt the membrane. The supernatant of OMVs was harvested by centrifugation at 12000 rpm for 1 min, boiled for 5 min in the Laemmli buffer, and applied for SDS-PAGE electrophoresis. The SDS-PAGE gel was visualized and recorded images under an excitation light of 488 nm before staining with the Coomassie brilliant blue.

**Cell culture and cytosolic protein delivery**

Hela, 293T, and CX-1 cells were cultured at 37°C with 5% $CO_2$ in Dulbecco's Modified Eagles Medium (DMEM, SIGMA) containing 10% Fetal Bovine Serum (FBS, HyClone) and 1% Penicillin Streptomycin solution. When reaching ~80% confluence, Hela, 293T, and CX-1 ($1 \times 10^4$ per well) cells were seeded into 24-well plate to incubate for 24 hours respectively. OMVs samples enriched with Dylight 488-labeled Spy or nanobodies were added into each well and incubated for 12h or 24h. For each cell line, the nuclei were stained with Hoechst 33342 (Beyotime Biotecnology), rinsed with PBS, fixed with 4% formaldehyde, and mounted with Antifade Mounting Medium. Cells incubated with OMVs were digested with trypsin-EDTA to produce monolayer cells, which were counted under the fluorescence microscope (OLYMPUS) to determine the transfection efficiency. To further analyze the uptake of OMVs by different cell lines *in vitro*, OMVs were labeled by a lipophilic membrane stain Dil fluorescent probe by co-incubation at 37°C for 20 min. The incubated OMVs were centrifuged and rinsed with PBS twice to remove free Dil fluorescent probes. Labeled

OMVs were then added to Hela, 293T, and CX-1 cells in 24-well plates for incubation. The incubated cells were rinsed with PBS and stained by Hoechst 33342 for imaging under a Nikon confocal laser scanning fluorescence microscope (CLSM) at the time point of 30 min, 6 h, and 24 h for each cell line respectively. To explore the mechanism of endocytosis of OMVs, Hela, 293 T, and CX-1 cells were cultured in 24-well plates. A final concentration of 80 μM Dynasore, the inhibitor of endocytosis, was added 30 min before the addition of the Dil labeled-OMVs. 6 h later, cells were checked by CLSM.[2]

**CCK-8 assay**

Hela, 293T, and CX-1 cells ($3\times10^3$per well) were inoculated in a 96-well plate and cultured overnight. OMVs (10μL) were added into each well, and PBS (10μL) was used as the control. CCK-8 was added to each well and incubated at 37°C for 1h. Samples were collected at 0h, 24h, 48h, and 72h and the absorbance at 450 nm was subsequently measured by a microplate reader (BioTek SYNERGY H1).

**Enzyme Linked Immunosorbent Assay (ELISA)**

ELISA was applied to detect the activity of proteins delivered into cells by OMVs. The sample of OMVs enriching Pfu nanobodies fused with His-tag was cultivated with Hela cells at 37°C for 24h. The cell culture was lysed with RIPA (Radio-Immunoprecipitation Assay, Solarbio) buffer, and the supernatant was collected by centrifugation twice at 12000 rpm for 5 min and was then added to the 96-well plate coated with 100 μL/well of 10ng Pfu protein, following by incubation at 37°C for 1h. Cell lysate without OMVs was used as the control. 10ng/ml Pfu nanobodies was used as positive control. The plate was washed 5 times with 250 μL PBST (PBS with Tween 20) and incubated with 100 μL anti-His antibody at the ratio of 1:5000 at 37°C for 1h. Then the plate was washed with 250 μL PBST 5 times and added with 50 μL streptavidin-HRP at the ratio of 1:5000 for further incubation at 37°C for 1h. Then the plate was washed with 250 μL PBST 5 times and subsequently incubated with 50 μL TMB (3, 3', 5, 5'-tetramethylbenzidine) substrate at 37°C for 15 min. The reaction was terminated by adding 50 μL concentrated sulfuric acid, and the signals were measured at the absorbance of 450 nm.

**Data analysis**

Data was collected at least three times and presented with mean ± standard deviation. Student's t-tests were employed to designate statistical significance.
